# Distinct roles of the polarity factors Boi1 and Boi2 in the control of exocytosis and abscission in budding yeast

**DOI:** 10.1101/151670

**Authors:** Aina Masgrau, Andrea Battola, Trinidad Sanmartin, Leszek P. Pryszcz, Toni Gabaldón, Manuel Mendoza

## Abstract

Boi1 and Boi2 (Boi1/2) are budding yeast plasma membrane proteins that function in polarized growth, and in cytokinesis inhibition in response to chromosome bridges via the NoCut abscission checkpoint. How Boi1/2 act in these two distinct processes is not understood. We demonstrate that Boi1/2 are required for a late step in the fusion of secretory vesicles with the plasma membrane of the growing bud. Cells lacking Boi1/2 accumulate secretory vesicles and are defective in bud growth. In contrast, Boi2 is specifically required for abscission inhibition in cells with chromatin bridges. The SH3 domain of Boi2, which is dispensable for bud growth and targets Boi2 to the site of abscission, is essential for abscission inhibition. Gain of function of the exocyst, a conserved protein complex involved in tethering of exocytic vesicles to the plasma membrane, rescued secretion and bud growth defects in *boi* mutant cells, and abrogated NoCut checkpoint function. Thus, Boi2 functions redundantly with Boi1 to promote the fusion of secretory vesicles with the plasma membrane at sites of polarized growth, and acts as an abscission inhibitor during cytokinesis in response to chromatin bridges.

## Introduction

Exocytosis, the delivery of secretory vesicles containing new membranes and membrane-remodelling factors to the plasma membrane (PM), is essential for cell growth and division. The molecular principles of exocytosis have been well characterized in the budding yeast *Saccharomyces cerevisiae*. In this organism, secretory vesicles are transported towards growth sites in the bud by actin-based transport; the actin cytoskeleton is in turn polarized by the localized activation of membrane-associated Cdc42 at the prospective bud site (Park and Bi, 2007). Secretory vesicles then fuse with the PM through soluble N-ethylmaleimide-sensitive factor attachment protein receptor (SNARE) complex formation. Prior to SNARE-mediated fusion with the target membrane, secretion requires a rate-limiting step known as “tethering”, mediated by an evolutionarily conserved octameric complex, the exocyst (Wu *et al.*, 2008; He and Guo, 2009). Exocyst function is essential for cell growth and its inactivation leads to accumulation of exocytic vesicles in the cytoplasm (Novick et al., 1980; Guo et al., 1999; He et al., 2007; Wu et al., 2010). The Sec3 and Exo70 subunits associate directly with the PM in a manner that is largely independent of the actin cytoskeleton, whereas the other six subunits (Sec5, 6, 8, 10, 15, and 84) are transported to growth sites on membrane vesicles. This has led to the hypothesis that assembly of all subunits at the plasma membrane mediates vesicle tethering prior to fusion (Boyd et al., 2004).

Exocytosis is also important for completion of cytokinesis. During this process, contraction of a membrane-associated actomyosin ring guides ingression of the PM at the site of cell division. Membrane resolution then splits the cell into two distinct topological units, in a process known as abscission (Green et al., 2012; Schiel and Prekeris, 2012). In HeLa cells, inactivation of the exocyst during cytokinesis leads to abscission defects (Gromley et al., 2005). In budding yeast, actomyosin ring contraction at the site of division, called the bud neck, is coupled to synthesis of a chitin-based primary septum (PS). Yeast exocyst mutants show aberrant ring contraction dynamics, mislocalization of the PS chitin synthase Chs2, and cytokinesis defects suggesting that exocytosis is also required for late cell division steps in this organism (Dobbelaere and Barral, 2004; VerPlank and Li, 2005). The timing of abscission is monitored by a mechanism known as NoCut in budding yeast, and the abscission checkpoint in animal cells, which inhibits abscission in cells with chromatin bridges caught in the path of the cell division machinery (Steigemann *et al.*, 2009; Nähse *et al.*, 2016; Amaral *et al.*, 2016a; 2016b). Whether NoCut impinges on the exocytic machinery is not known.

The functionally redundant yeast cortical proteins Boi1 and Boi2 (Boi1/2) were previously implicated in polarized growth (Bender et al., 1996; Matsui et al., 1996) and in the NoCut abscission checkpoint (Norden et al., 2006; Mendoza et al., 2009). How Boi1 and Boi2 act in these two distinct processes remained unclear. Here, we demonstrate that Boi1/2 promote vesicle exocytosis during bud growth. In contrast, the SH3 domain of Boi2, which targets the protein to the bud neck, is dispensable for bud growth but is specifically required to inhibit abscission in cells with chromatin bridges. Our results raise the possibility that Boi2 acts as both an activator of exocytosis and an inhibitor of abscission, depending on the cell cycle stage and the cellular response to chromosome segregation defects during cytokinesis.

## Results

### Boi1 and Boi2 are essential for bud growth

Single *boi1Δ* and *boi2Δ* cells show normal growth and morphology but double deletion mutants display severe morphogenesis and growth defects (Bender et al., 1996; Matsui et al., 1996) (Fig. **S1A**). To investigate the role of Boi1/2 during cell growth, we generated a conditional *boi1 boi2* null strain using an auxin-inducible degron (AID) to rapidly target Boi2 for poly-ubiquitination and proteasome-dependent degradation in the presence of 1-naphthaleneacetic acid (NAA) and the plant E2 ligase Tir1 (Nishimura et al., 2009) (Fig. **1A**). A *boi2-aid* strain expressing Tir1 grew well in complete media but failed to form colonies in the presence of NAA specifically in a *boi1Δ* background, consistent with an essential role of Boi1/2 in cell viability (Fig. **1B**).

**Figure 1.**
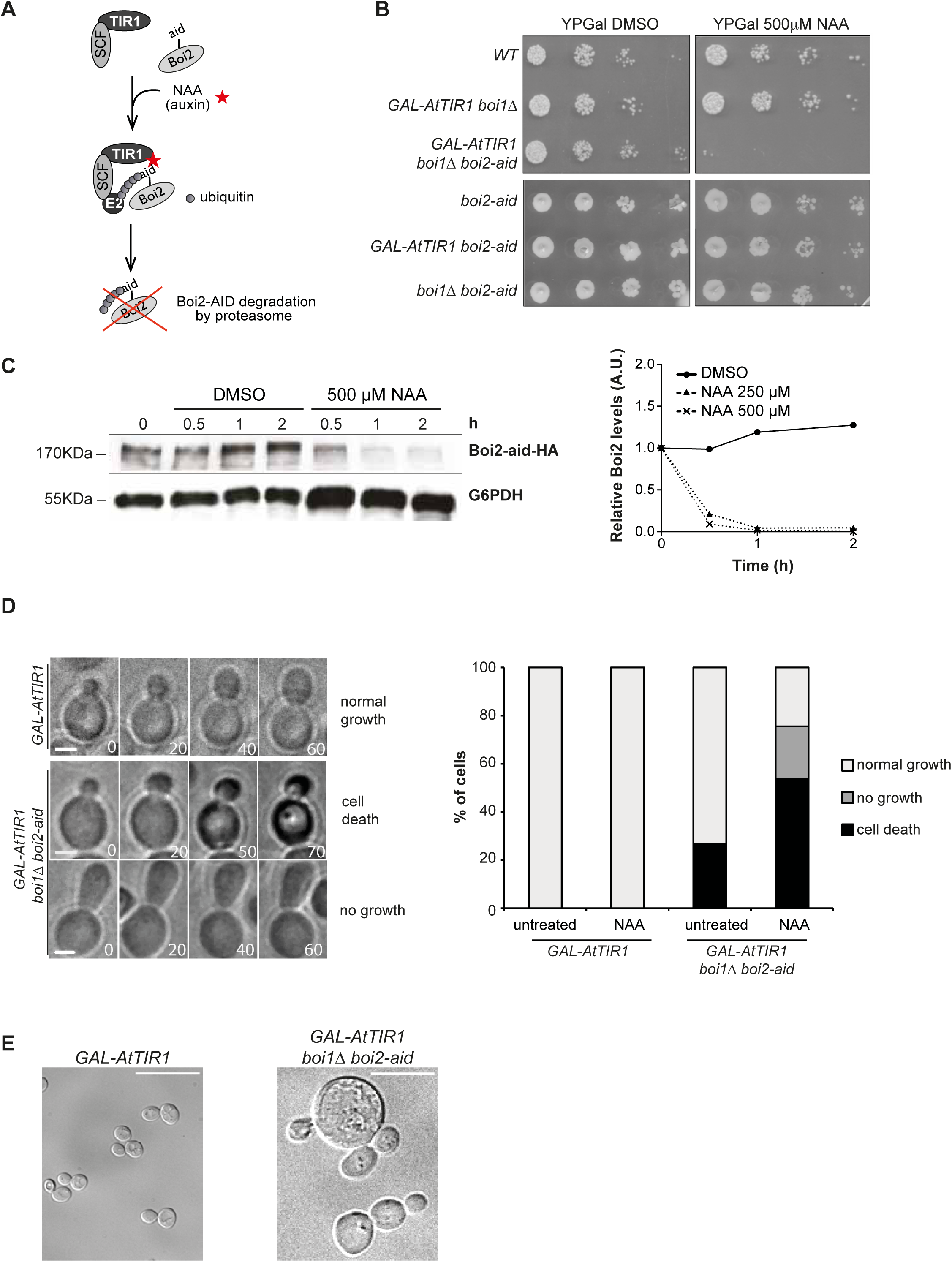
Boi1 and Boi2 are essential for bud growth. (A) Schematic representation of the auxin-induced degron system. SCF: Skp1, Cullin, and F-box complex. E2: E2 ubiquitin ligase. Aid: auxin-inducible degron. (B) Serial dilutions of the indicated strains spotted onto the indicated plates and incubated for 3 days at 30 °C. (C) *GAL-AtTIR1 boi1Δ boi2-aid-HA* cultures were grown in YPR to log phase, transferred to YPG for 2 h, and Boi2-aid-HA was detected by immunoblotting at the indicated time points after addition of DMSO or the indicated concentrations of NAA. Glucose-6-phosphate dehydrogenase (G6PDH) was used as a loading control. (D) DIC time-lapse imaging of wild type and *boi1Δ boi2-aid* mutants expressing *GAL-AtTIR1*. Cells were grown in YPR to log phase, transferred to YPG for 2 h, and imaged in the presence of 250 μM NAA for the following 6 hours. Cell images (4 μm thick stacks spaced 0.8 μm) were acquired every 5 min; selected frames are shown. Time is represented in minutes. Scale bars, 2 μm. Cells were classified in three groups as indicated; the relative frequencies of these groups are indicated in the graph. N > 23 cells pooled from 2 independent experiments. (E) DIC images of wild type and *boi1Δ boi2-aid* cells 24 h after addition of NAA. Scale bar, 10 μm.

To determine the consequences of Boi1/2 depletion on cell growth, wild type and *boi1Δ boi2-aid* cells were examined by differential interference contrast (DIC) time-lapse microscopy 2 h after addition of 0.25 mM NAA, when Boi2-aid protein levels were reduced to nearly undetectable levels (Fig. **1C**). Wild type and *boi1Δ boi2-aid* cells had similar morphology; however, Boi-depleted cells were severely impaired in surface growth. NAA-treated *boi1Δ boi2-aid* cells with small or medium buds grew at a slower rate or turned dark and stopped growing altogether (Fig. **1D**). Moreover, large round *boi1Δ boi2-aid* cells were observed 24 hours after NAA addition (Fig. **1E**). Thus Boi1/2 function in cell growth, and depolarized growth previously reported in *boi1Δ boi2Δ* mutants (Bender et al., 1996; Matsui et al., 1996) might represent secondary defects associated with long-term Boi1/2 depletion.

### Boi1/2 are dispensable for organization of actin and cell polarity factors

To assess if Boi1/2 are required for maintenance of a polarized actin cytoskeleton, F-actin was visualized with fluorescently labelled phalloidin in wild type and *boi1Δ boi2-aid* cells treated with NAA for 2 h. Actin patches and cables appeared similarly organized in wild type and Boi-depleted cells (Fig. **2A**). In addition, we determined the localization of various cell polarity proteins fused to GFP in wild type and Boi-depleted cells. Lack of Boi1/2 did not severely affect the localization of the Cdc42 guanine nucleotide exchange factor Cdc24, whereas it did moderately reduce that of the Boi-interacting protein Bem1 (Mccusker et al., 2007) and of the exocyst components Sec3, Exo70 and Exo84 to the bud cortex. Furthermore, Boi-depleted cells showed slightly increased frequency of bud cortex localization of the Rab-like protein residing on post-Golgi vesicles, Sec4 (Goud et al., 1988) (Fig. **2B**). Thus Boi1/2 are not required for the maintenance of a polarized actin cytoskeleton and do not play a major role in the localization of polarity factors and exocytic vesicles to sites of polarized growth.

**Figure 2.**
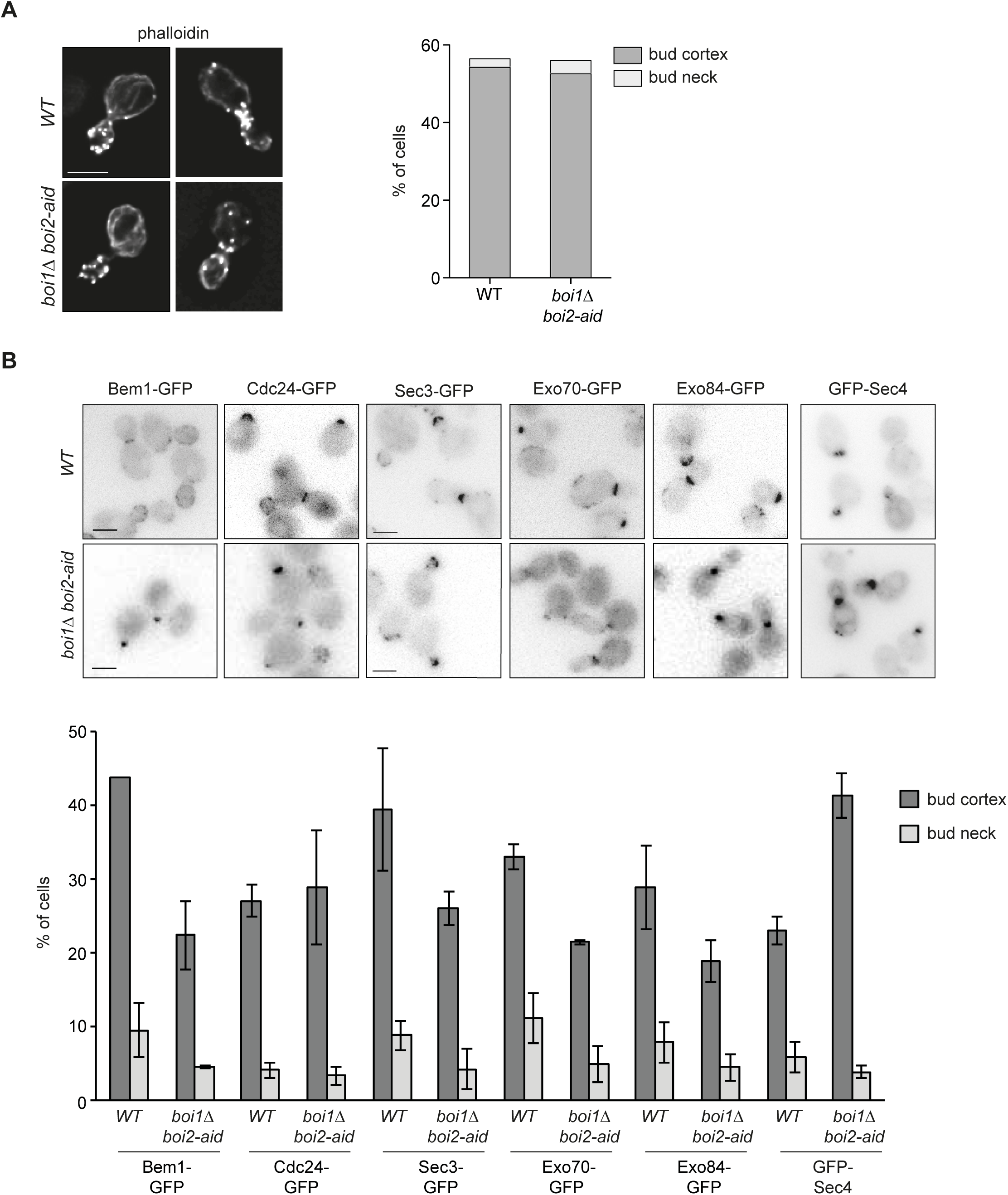
Boi1/2 are dispensable for actin organization or localization of cell polarity markers. (A) Distribution of filamentous actin (Alexa-488 phalloidin) in wild type and *GAL-AtTIR1 boi1Δ boi2-aid* cells 2h after NAA addition, treated as in Fig. 1D (N > 100). Scale bars, 5 μm. (B) Localization of GFP fusion proteins in wild type and *GAL-AtTIR1 boi1Δ boi2-aid* cells treated as in Fig. 1D. All fusion proteins were expressed from their chromosomal locus except GFP-Sec4, which was expressed from a centromeric plasmid. Results are represented as mean and SEM (N > 150 cells for each condition, two to four independent experiments).

### Identification of putative *boi1 boi2* suppressors by genome sequencing

To gain insight into the molecular functions of Boi1 and Boi2, we took advantage of a previously described *boi1Δ boi2Δ* strain, which was viable. These cells show no obvious morphological defects but are defective in the NoCut abscission checkpoint, which inhibits completion of cytokinesis in the presence of chromosome segregation defects (Norden et al., 2006; Mendoza et al., 2009). We hypothesized that this strain might contain one or more suppressor mutations (*SUP*) ensuring cell viability in the absence of Boi1/2. To identify putative suppressors, whole genome sequencing of this viable *boi1Δ boi2Δ* strain was performed and compared to its wild-type parent.

Sequence analysis showed that gene order and copy number were identical between the two strains, ruling out aneuploidy and gross genome rearrangements in *boi1Δ boi2Δ SUP* (see Materials and Methods, and Fig. **S2**). However, assessment of variation at the single nucleotide level identified 19 single nucleotide polymorphisms (SNPs) between *boi1Δ boi2Δ SUP* and its parental strain. Of these, 7 were predicted to introduce amino acid changes in the encoded proteins (see **Table S1**).

To determine linkage of SNPs to survival of *boi1Δ boi2Δ SUP* cells, genetic crosses were performed between this mutant and a *BOI1 BOI2* strain. As expected, a fraction of *boi1Δ boi2Δ* spores from this cross gave raise to viable colonies (Fig. **S1B**). One *boi1 boi2 SUP* clone was selected and backcrossed four more times; a *BOI1/boi1Δ BOI2*/*boi2Δ SUP/+* zygote that produced four viable spores was identified after the 5th backcross and its meiotic products were characterized by whole genome sequencing. Candidate suppressors should be present in both viable *boi1Δ boi2Δ* spores, and absent in the two *BOI1 BOI2* clones; SNPs fulfilling these criteria are shown in **Table S2**. Several SNPs clustered around specific genomic regions as expected by a selection sweep effect, presumably due to genetic background differences between *boi1Δ boi2Δ SUP* and the *BOI1 BOI2* strain (see Materials and Methods). However, the only SNP common to both the original and the backcrossed *boi1Δ boi2Δ SUP* segregants was located in the gene encoding the exocyst subunit Exo70, where it introduces one amino acid substitution (*EXO70-G388R*). Notably, the same mutation was independently isolated as a suppressor of *cdc42* and *rho3* mutants. The mutant protein, hereafter termed Exo70*, is expressed at a similar level to the wild-type protein and is incorporated into endogenous exocyst complexes, where it acts as a dominant, gain-of-function allele that suppresses the lethality and exocytosis defects of *cdc42* and *rho3* mutants (Wu et al., 2010).

### Boi1/2 are required for vesicle exocytosis

To test whether *EXO70\** is a dominant suppressor of the *boi1 boi2* mutant, centromeric plasmids encoding either the wild type or the dominant version of *EXO70* under the control of the regulatable *GAL1,10* promoter were introduced in *boi1Δ boi2-aid* strains. Galactose-driven expression of Exo70*\** was sufficient to restore growth of Boi-deficient cells in the presence of NAA, whereas over-expression of wild-type Exo70 did not (Fig. **3A**). *EXO70*\* expressed from its natural promoter at the endogenous locus supported growth of both *boi1Δ boi2-aid* cells in NAA-containing media (Fig. **3B**) and of *boi* null strains, although *boi1Δ boi2Δ EXO70\** strains grew at slower rates than wild type (Fig. **S1C**). Quantification of bud growth by DIC time-lapse microscopy showed that *boi1Δ boi2-aid EXO70\** cells grew in a polarized manner and at slower rates than wild type cells (Fig. **3C**). Together, these results indicate that Boi1/2 promote exocyst-dependent vesicle fusion with the PM during bud growth.

**Figure 3.**
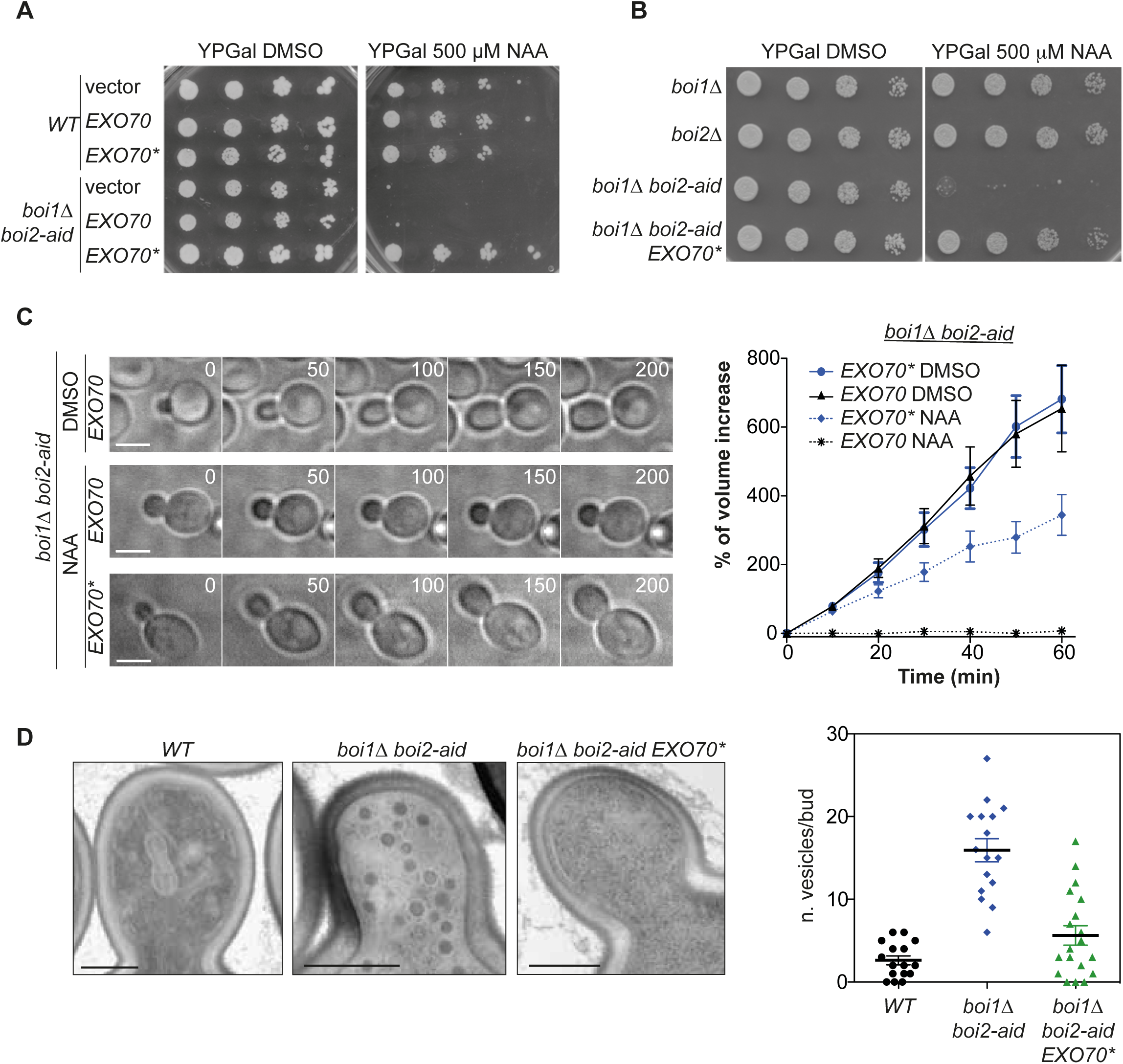
Boi1/2 are required for vesicle exocytosis. (A) Serial dilutions of wild type or *boi1Δ boi2-aid* strains bearing *EXO70, EXO70\** or empty centromeric (CEN) plasmids were grown on galactose-containing media with and without NAA for 4 days. The plasmid-borne *EXO70* gene was under the control of the GAL1 promoter. (B) Serial dilutions of cells of the indicated strains grown on YPG with and without NAA for 3 days. *EXO70\** is expressed from the *EXO70* promoter at the endogenous locus. (C) DIC time-lapse images of *boi1Δ boi2-aid* cells transformed with *EXO70* or *EXO70\** CEN plasmids. Cells were grown in SC Gal-Leu to log phase, and imaged 2 h after addition of 250 μM NAA at 25 °C. Numbers indicate time in minutes; scale bar, 5 μm. The graph represents the relative increase in volume of individual buds over time. N = 10 cells per condition. Error bars indicate SEM. (D) EM images of small budded cells in the indicated mutants. Cells were treated as in Fig. 1C and fixed 2 h after NAA addition. Scale bar, 500 nm. The number of vesicles/cell/section is represented in the graph (N> 15). Lines represent the mean, bars are SEM.

To directly assess this, electron microscopy (EM) analysis was performed in wild type and Boi-depleted cells. In wild type cells few secretory vesicles are detected by EM due to high basal secretion rates. In contrast, a higher number of 80-100 nm diameter vesicles were visualized in NAA-treated *boi1Δ boi2-aid* cells (Fig. **3D**). Secretory vesicles were particularly abundant near the surface of small and medium-sized buds, in a manner reminiscent of exocyst loss-of-function mutants. Conversely, vesicle accumulation was largely alleviated *in boi1Δ boi2-aid EXO70*\* cells (Fig. **3D**). Thus, Boi1 and Boi2 may promote the tethering function of the exocyst to allow secretory vesicles to fuse with the plasma membrane.

We then tested whether Boi1 and Boi2 are required for the secretion of specific protein cargoes. Budding yeast cells have two major types of exocytic vesicles, which can be distinguished by their density and protein content: light vesicles containing cell wall-remodelling cargo like the endoglucanase Bgl2, and dense vesicles typified by the presence of invertase (Harsay and Bretscher, 1995). The activity of secreted periplasmic invertase in wild type and Boi-depleted cells was determined using a standard assay (Goldstein and Lampen, 1975). Exocyst mutant cells (*sec6-4*) known to be impaired in secretion of invertase showed a strong reduction in the activity of the secreted form of this enzyme at the restrictive temperature, relative to wild type cells (He et al., 2007) (Fig. **4A**). In contrast, *boi1Δ boi2-aid* cells showed no reduction of secreted invertase activity after incubation with NAA to deplete Boi2, suggesting that Boi1/2 are not required for secretion of invertase-containing vesicles (Fig. **4A**). To assay secretion of Bgl2, immunoblots were used to probe the internal fraction of the enzyme in spheroplasts of wild type and mutant cells. Both *sec14-1* cells previously reported to be blocked in secretion of Bgl2 at 37 °C (Curwin et al., 2009), and Boi-depleted cells accumulated large amounts of Bgl2 in the cytoplasm relative to the total cellular content. This defect was fully reversed in the *boi1Δ boi2-aid EXO70\** strain (Fig. **4B** and **S3**). Thus, Boi1/2 function is dispensable for secretion of invertase vesicles but is essential for exocytosis of Bgl2-type vesicles, and this requirement can be bypassed by gain-of-function of the exocyst.

**Figure 4.**
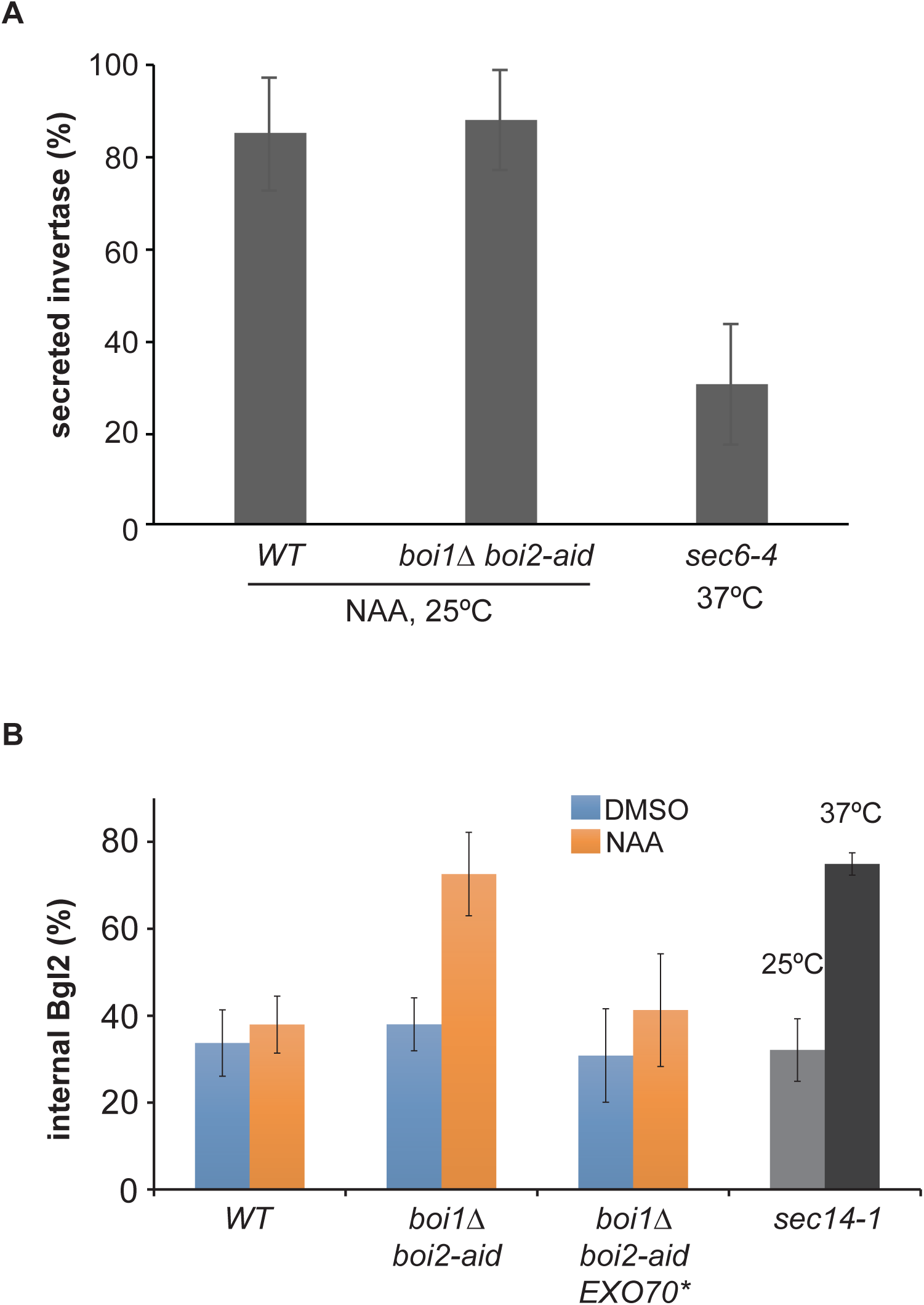
Boi1/2 are essential for secretion of Bgl2 but not invertase. (A) Activity of secreted invertase in wild type and *boi1Δ boi2-aid* cells treated with NAA for 2 h, and of *sec6-4* cells shifted to 37 °C for 2 h. Invertase secretion is the ratio of external invertase activity relative to total (internal + external) activity. Invertase data are the mean and SEM of five independent experiments performed in triplicate. (B) Internal Bgl2 levels of cells of the indicated strains after incubation with DMSO or NAA for 2 h (*boi1Δ boi2-aid*), or after shift to 37 °C for 2 h (*sec14-1*). Bgl2 levels were determined by western blot and represent the ratio between internal and total (internal + external). G6PDH was used as a loading control. Bgl2 data are the mean and SD of three independent experiments.

### Boi2 inhibits cytokinesis in cells with catenated chromatin bridges

We next addressed the role of Boi1/2 in cytokinesis. Anaphase chromatin bridges lead to inhibition of abscission; this inhibition was bypassed in *boi1Δ boi2Δ* cells, leading to the conclusion that Boi1/2 function at the bud neck as abscission inhibitors (Norden et al., 2006; Mendoza et al., 2009). Our finding that Boi-deficient cells used in these studies contained the *EXO70*\* suppressor is in line with their lack of growth defects, but raised the possibility that their failure to inhibit abscission could be partially or even completely due to exocyst gain of function, and not to loss of Boi1/2. To address these issues, we revisited the role of Boi1/2 in cytokinesis. Wild-type, *boi1Δ* and *boi2Δ* cells were released from a G1 arrest and shifted to 37 °C to induce chromatin bridges by inactivation of topoisomerase II with the *top2-4* mutation (Holm et al., 1985). The spindle pole body (SPB) marker Spc42-GFP was used to monitor progression through mitosis, whereas contraction and subsequent resolution of the PM at the bud neck were monitored using GFP fused to the PM targeting CAAX motif of Ras2 (Amaral *et al.*, 2016a).

Live fluorescence microscopy showed that wild type cells underwent ingression of the plasma membrane at the bud neck after entry of the SPB in the bud, and this was followed by membrane resolution into two distinct layers as previously reported (abscission) (Fig. **5A-B**). In *boi1Δ* and *boi2Δ* cells, ingressed bud neck membranes resolved with dynamics similar to those of wild type cells (Fig. **5B**). In contrast, *top2-4* and *top2-4 boi1Δ* mutants showed ingression of the PM at the bud neck but were impaired or severely delayed in its resolution, consistent with a defect in abscission in *top2* mutants (Fig. **5A-B**) as previously observed by fluorescence microscopy and EM tomography (Amaral *et al.*, 2016a). However, analysis of GFP-CAAX showed that unlike *top2-4* and *top2-4 boi1Δ* mutants, most *top2-4 boi2Δ* cells were able to complete abscission (Fig. **5A-B**). Therefore, Boi2 is specifically required for inhibition of abscission in cells with catenated chromatin bridges, whereas Boi1 is dispensable for this inhibition.

**Figure 5.**
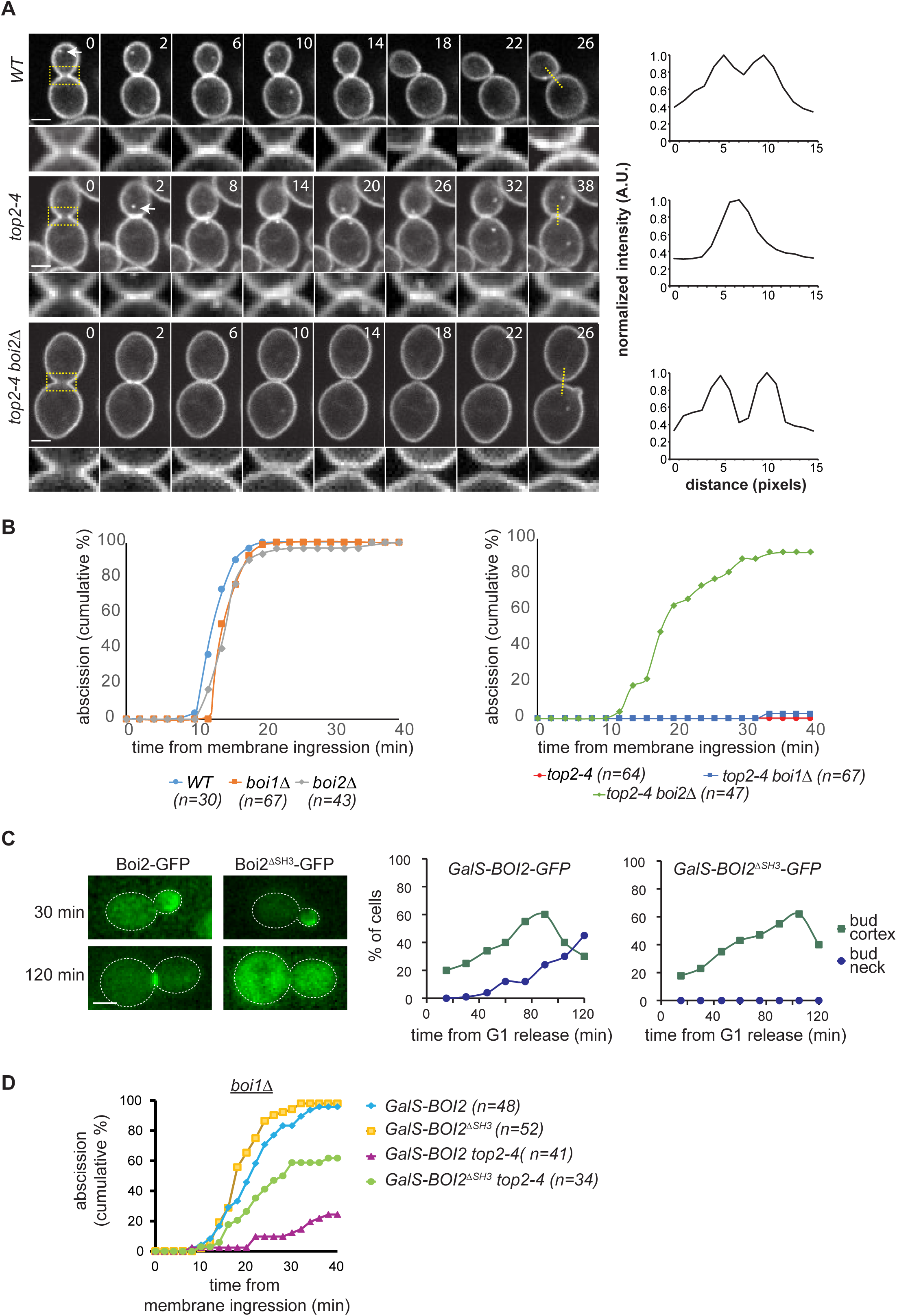
Boi2 inhibits cytokinesis in *top2-4* cells through its SH3 domain. (A) Membrane ingression and abscission in representative cells of the indicated strains. Z-stacks spaced 0.3 *μ* m apart and spanning the whole cell were acquired at 2-min intervals, but only central Z-planes are shown. Whole-cell images are shown on top, and enlargements of the bud neck region at the bottom. The spindle pole marker Spc42-GFP (visible only in some Z-sections; arrows) allowed the simultaneous visualization of spindle elongation. Numbers indicate time in minutes; time 0 marks the frame before membrane ingression. To analyze the status of the bud neck membrane, the GFP fluorescence intensity was measured across the cleavage plane in the central Z-plane (yellow lines) shown in the right. A reduction in local intensity marked membrane resolution and was scored as abscission (top and bottom graphs); a single peak denoted the pre-abscission stage (middle). (B) Graphs show the fraction of cells completing abscission relative to the time of membrane ingression. Cells were synchronized in G1 with alpha-factor in YPD at 25°C and shifted to 37°C 15 minutes before release from the G1 block. Data are from cells pooled from 2-4 independent experiments. (C) Cells of the indicated strains were released from a G1 block and imaged at time intervals in galactose-containing media. The localization of Boi2 to either the cortex of the bud neck (left) was determined by wide-field microscopy. At least 30 cells were analysed per time point. (D) Abscission analysis as in (A-B), except that the cultures were grown to mid-log phase in galactose-containing media. Data are from cells pooled from 2-3 independent experiments.

Boi1 and Boi2 share a similar domain organization, featuring a Pleckstrin-homology (PH) domain at their C-terminus and a Src-homology 3 (SH3) domain at their N-terminus. Boi1/2 PH domains can interact with PM lipids and associate with the bud cortex; moreover, the PH domain of Boi1 is required for viability of *boi1 boi2* mutant cells (Hallett *et al.*, 2002; Yu *et al.*, 2004). Therefore, PH domains might mediate the essential function of Boi1/2 in bud growth. In contrast, the SH3 domain of Boi1 is not required for viability of *boi* null mutants and targets Boi1 to the bud neck (Hallett *et al.*, 2002). We therefore asked whether the SH3 domain of Boi2 might be required for its bud neck localization and its function in abscission inhibition. A version of Boi2 expressed under the control of the weak GalS promoter, and lacking the 102 N-terminal aminoacids spanning the SH3 domain, did not perturb Boi2 targeting to the bud cortex, but abrogated its localization to the bud neck during cytokinesis (Fig. **5C**). This Boi2 mutant version (Boi2^ΔSH3^) also supported normal bud growth and abscission in *boi1Δ* cells (Fig. **5D**). However, *boi1Δ GalS-boi2^ΔSH3^* cells did not inhibit abscission in the presence of chromatin bridges caused by Topoisomerase II inactivation (Fig. **5D**). Thus, the SH3 domain of Boi2 is not essential for bud growth, but is specifically required for Boi2 targeting to the bud neck and for its function in the NoCut abscission checkpoint.

### Exocyst gain of function restores abscission in cells with chromatin bridges

As inhibition of the exocyst perturbs the completion of yeast cytokinesis (Dobbelaere and Barral, 2004; VerPlank and Li, 2005), we asked whether gain of function of the exocyst can prevent its inhibition. To this end, the GFP-CAAX reporter was used to monitor abscission in *EXO70*\* cells with normal and defective chromosome segregation. *EXO70*\* cells completed abscission with similar kinetics to those of wild type, and importantly, failed to inhibit abscission in the presence of catenated DNA bridges (Fig. **6A**). These results raise the possibility that abscission inhibition in cells with chromatin bridges requires modulation of exocyst-dependent, membrane-remodelling events at the abscission site. To gain insight into cytokinesis membrane trafficking events in cells with chromatin bridges, we followed the kinetics of the vesicle marker Sec4, which accumulates in the bud neck at the end of mitosis and controls membrane trafficking during cytokinesis (Lepore *et al.*, 2016) (Fig. **6B**). The levels of Sec4-GFP accumulating in the bud neck relative to total Sec4 cellular levels were significantly higher in *top2-4* cells than in wild type cells. Moreover, increased Sec4 levels at the bud neck in the presence of chromatin bridges were restored in *EXO70\** cells but not in *boi2Δ* cells (Fig. **6B-C** and **S4**). Thus, cytokinetic membrane remodelling is altered in response to DNA decatenation defects, in a process that is independent from Boi2 but can be bypassed by gain of function of the exocyst.

**Figure 6.**
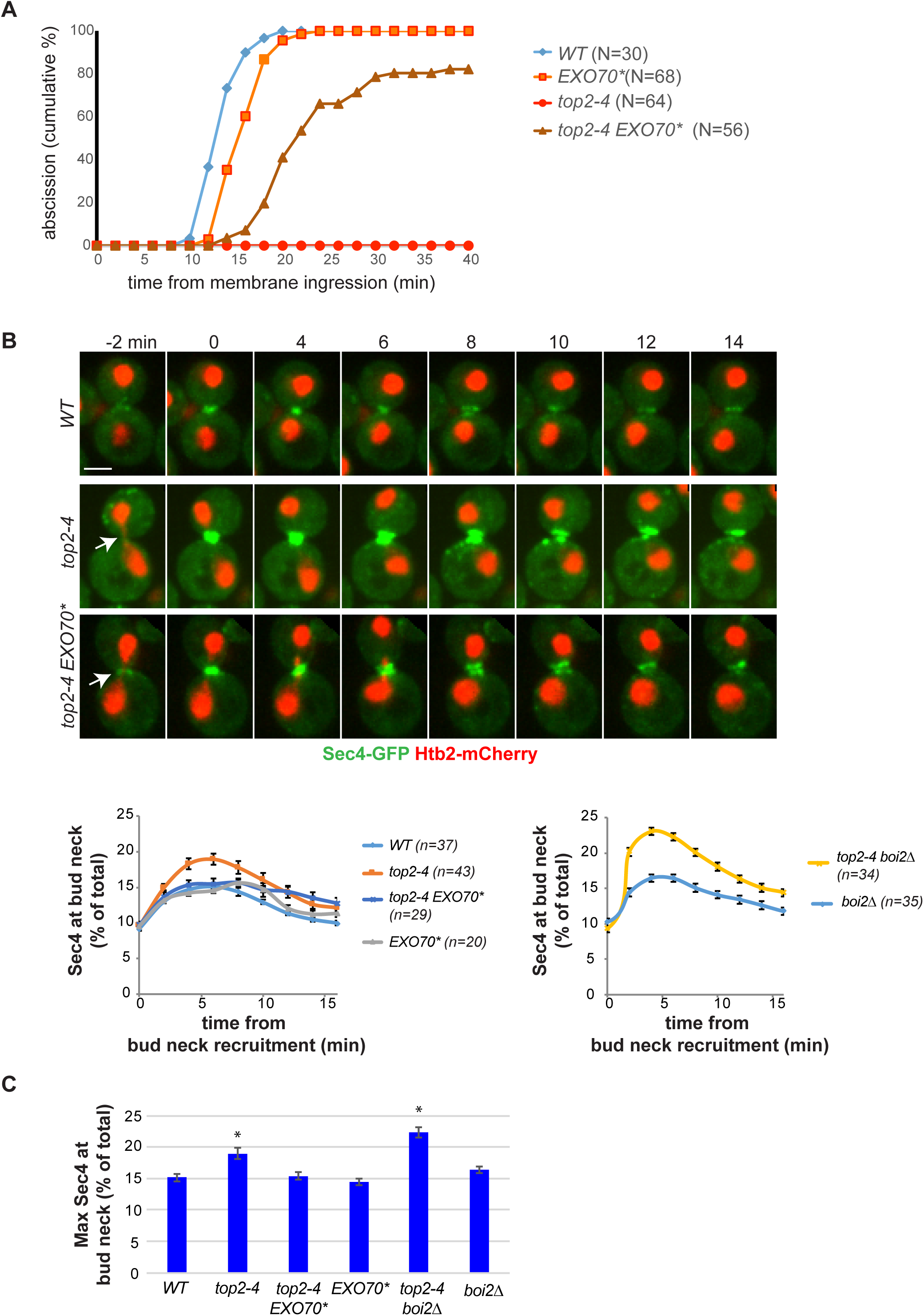
*EXO70*\* is a suppressor of abscission defects in topoisomerase II-deficient cells. (A) Abscission analysis as in Fig. 5A-B. Data are from cells pooled from 2-3 independent experiments. (B) Localization of Sec4-GFP and histone 2B (Htb2) fused to mCherry in late mitotic cells. Arrows mark chromatin bridges in *top2-4* mutants. Time 0 marks the appearance of Sec4 at the bud neck. Levels of Sec4-GFP at the bud neck, expressed as % of the total GFP fluorescence in the cell, were measured in sum projections of confocal z-stacks spanning the entire cell. Mean and SEM are shown. Data are from cells pooled from 2-4 experiments. (C) Maximal levels (mean and SEM) of Sec4-GFP at the bud neck of the indicated strains. Asterisks denote statistically significant differences from the wild type (p<0.05, Student’s t-test).

## Discussion

In this study, we have shown that Boi1/2 are required specifically for exocytosis of specific vesicle types and for bud growth, whereas Boi2 inhibits abscission in response to DNA decatenation defects. Our results suggest that Boi1/2 function in secretion of Bgl2-containing vesicles, but not of invertase vesicles. Most exocyst mutants (including *exo70* alleles) are blocked in secretion of both vesicle types (Novick et al., 1980; Harsay and Bretscher, 1995; Guo et al., 1999; Wu et al., 2010). However the hypomorphic *exo70-35* and *exo70-38* mutants, like Boi1/2-depleted cells, are specifically impaired in secretion of Bgl2 but not invertase vesicles (He et al., 2007). Boi1 and Boi2 interact with the polarity factor Bem1 in vivo, and with exocyst components in vitro and by two-hybrid assays (Tonikian et al., 2009); moreover, Boi1/2 and these other cell polarity factors associate with the plasma membrane of the growing bud (Hallett et al., 2002; Yu et al., 2004). Therefore, Boi1/2 might specifically target Bgl2 (and perhaps other) vesicles for tethering by exocyst complexes during bud growth.

In both yeast and human cells, chromatin bridges can cause inhibition of abscission through the NoCut abscission checkpoint, but the molecular mechanisms involved remain poorly understood. In human cells, this process involves regulation of the Endosomal Sorting Complex Required for Transport (ESCRT) III complex, which associates with the plasma membrane at the site of cytokinesis and regulates abscission timing (Carlton et al., 2012). Our data indicates that in yeast, abscission inhibition relies on association of Boi2 with the abscission site. Although both Boi-dependent functions, in polarized growth and abscission, may involve plasma membrane remodelling processes, the relevant molecular mechanisms are probably distinct. Indeed, the essential role of Boi1 and Boi2 for exocytosis and polarized growth probably depends on their PH domains, which are required for association with the bud cortex and are essential for cell viability (Hallett *et al.*, 2002). In contrast, we find that the SH3 domain of Boi2 is dispensable for viability, but is essential for bud neck targeting and inhibition of abscission. Interestingly, the SH3 domains of Boi1 and Boi2 can interact with multiple proteins involved in polarized growth, including exocyst components (Tonikian *et al.*, 2009). Whether these interactions are relevant for the control of abscission in cells with chromosome segregation defects remains an important question for future studies.

Finally, the finding that gain of function of the exocyst can restore abscission in yeast cells with chromatin bridges open the possibility that exocyst regulation may play a role in the inhibition of abscission in yeast, and perhaps also in human cells. We note that although the accumulation of Sec4-GFP at the abscission site in *top2-ts* cells may represent inhibition of exocytosis during cytokinesis, EM imaging failed to detect vesicles in the vicinity of the bud neck during cytokinesis in *top2* mutants [(Amaral 2016a) and data not shown]. Alternatively, accumulation of Sec4 in the bud neck of *top2* mutant cells may reflect alterations in a non-exocytic membrane-remodelling process such as endocytosis, which can also be regulated by the exocyst (Jose *et al.*, 2015). A detailed characterization of specific membrane and protein trafficking events during abscission, and the potential role of exocyst complexes in regulating these processes, should provide valuable insight into the mechanisms of abscission regulation in response to chromatin bridges.

## Materials and methods

### Strains and media

*S. cerevisiae* strains are derivatives of S288c, except *boi1Δ boi2Δ* and its parental wild type strain (kind gifts from E. Bailly, INSERM) which have the BFA264-15D background. Yeast cells were grown in YPD/YPG/YPR (1% bacto-yeast extract; 2% bacto-peptone; 2% dextrose, galactose or raffinose; and 0.004% adenine). Gene tagging and deletions were generated by standard PCR-based methods. The *EXO70\** (G488R) plasmid was generated by site-directed mutagenesis (Quickchange, Stratagene) of a *GAL1-EXO70* centromeric vector (Open Biosystems). To integrate *EXO70\** in its native locus in S288c strains, *EXO70\** was tagged with the HAnatNT2 cassette (Janke et al., 2004) in *boi1Δ boi2Δ SUP*, and a PCR-derived fragment of *EXO70\**-*HA*:*natNT2* was used in a one-step allele replacement; correct strains were identified by sequencing.

### Time-lapse and Fluorescence microscopy

Time-lapse microscopy was performed on cells in log phase, or after synchronization with alpha factor as indicated, plated in minimal synthetic medium in concanavalin A-coated Lab-Tek chambers (Nunc) and placed in a pre-equilibrated temperature-controlled chamber. Imaging was performed using a Cell Observer HS microscope with a 100x, 1.4 NA objective and an AxioCam MrX camera (Zeiss) (Fig. 1 and 3), an AF6000 wide-field microscope (Leica) with a 100x, 1.4 NA objective and an iXon 885 DU EM-CCD camera (Andor; Fig. 2 and 5C), or a Revolution XD spinning disc confocal microscope with a 100x, 1.45 NA objective and an iXon 897E Dual Mode EM-CCD camera (Andor; Fig. 5, 6 and S4). Bud volumes were calculated from DIC stacks using ImageJ and the BudJ plug-in (Ferrezuelo et al., 2012) customized for analysis of Z-stacks. To visualize F-actin, cells were fixed with 3.7% formaldehyde for 1 h, washed in PBS, incubated with 0.2 U/ml Alexa 488-phalloidin for 1 h at 25 °C, washed in PBS and visualized immediately in 80% glycerol / 20% PBS. Abscission assays were performed as in (Amaral *et al.*, 2016a). Briefly, the plasma membrane was visualized via *GAL1*,*10* promoter driven GFP-CAAX integrated in the *HIS3* locus. Expression of GFP-CAAX in glucose media was driven by the chimeric *ADH1pr*-Gal4-ER-VP16 transcription factor (Louvion et al., 1993) (*URA3*; gift of Francesc Posas, UPF) and addition of 90 nM β-estradiol (Sigma) 2 h before imaging. Only cells starting cytokinesis (membrane ingression) at least 40 min before the end of image acquisition were considered for the quantifications of abscission. Fluorescence intensities were measured from single sections of each cell to score membrane separation (abscission).

### Electron microscopy

Cells were cryoimmobilized using a EM HPM 100 high-pressure freezer (Leica), freeze-substituted in anhydrous acetone containing 2% glutaraldehyde and 0.1% uranyl acetate (Fig. 3) or 2% OsO4 and 0.1% uranyl acetate (Fig. 5) and warmed to room temperature (EM AFS-2, Leica). After several acetone rinses, cells were incubated with 1% tannic acid, washed, incubated with OsO4 at 1% acetone, washed again, infiltrated with Epon resin and embedded and polymerised at 60 °C. Ultrathin sections were obtained using an Ultracut UC6 ultramicrotome (Leica) and observed in a Tecnai Spirit electron microscope (FEI Company, The Netherlands) equipped with a Megaview III CCD camera.

### Sequencing and bioinformatics analysis

Strains were sequenced using Illumina Genome Analyzer II or HiSeq paired-end technology. Reads were trimmed before the genome assembly at the base with quality lower than 20 using FASTX-Toolkit (Cold Spring Harbor Laboratory, http://hannonlab.cshl.edu/fastx_toolkit). Subsequently, reads shorter than 31 bases (and their pairs) were discarded. Assemblies for each strain were created de novo using Velvet v. 1.1.02 (Zerbino and Birney, 2008). The insert size of the paired-end reads was estimated by aligning reads onto the assembly created from single reads. The optimal k-mer length (k) for each strain was chosen using VelvetOptmiser.pl to maximize the total number of bases in contigs longer than 1 kb (Lbp). In addition, we used auto-estimation of coverage cut-off and removed contigs shorter than 1 kb. Table S3 lists detailed information about each resulting assembly. Genomic reads were aligned onto the SGD reference strain using Bowtie2 v. 2.0.0 - beta7 (Langmead and Salzberg, 2012). SNPs were detected using bam2snp.ref.py v1.0. In brief, SNPs were called at sites with different genotypes in a given strain than in the wild-type strain. Only SNPs with high coverage (≥ 10x) and low ambiguity (≥ 90% of aligned reads call the same nucleotide at that position) at both the mapped reads of the given mutant strain and the reference were accepted. This filtering procedure was able to discard likely false-positive SNPs.

Depth-of-coverage variation was analyzed with IGV v 2.3.32 (Thorvaldsdóttir et al., 2013) to identify potential duplications or deletions. In addition, we analyzed read pairs with incongruent distance or orientation in order to identify possible rearrangements ie. inversions, deletions, duplications or translocations using bam2sv.py v1.0b. Finally, we used de novo contigs aligned against wild-type reference assembly as an independent line of evidence for potential rearrangements detection. Alignments were created by MUMer v. 3.07 with default parameters (Kurtz et al., 2004). Only best reciprocal matches were further accepted, but partial overlaps between matches were allowed. A minimal alignment length of 65 bp was set as a threshold. In addition, cut-off of 200 bp was set for deletion detection.

Genomic libraries were deposited to Short Read Archive (PRJEB7711). bam2sv.py and bam2snp.ref.py are available at https://bitbucket.org/lpryszcz/bin.

### Secretion assays

Invertase secretion was assessed essentially as described previously (Goldstein and Lampen, 1975). Briefly, cells were grown to logarithmic phase and transferred to 0.1% glucose YPDA to induce invertase expression, either in the presence of 90 nM β-estradiol and 0.5 mM NAA (to respectively induce Gal1-*At*Tir1 and Boi2 degradation in *boi1Δ boi2-aid*) or at 37 °C (for *sec6-4*). After 2 hours, cells were centrifuged and washed with cold 10 mM NaN_3_, and equal cell numbers were transferred to tubes with 10 mM NaN_3_ / 0.2% Triton-X100, which then were vortexed and freeze-thawed (total invertase pool), and 10 mM NaN_3_ (external pool). In triplicate, adjusted amounts of sample were diluted in 0.1 M sodium acetate (pH 5.1) and 0.5 M sucrose was added at timed intervals to measure invertase activity. Reactions were stopped after 30 min with 0.2 M KPO_4_ pH 7 / 10 mM Nethylmaleimide, and boiled. Glucose was detected with GO Assay Kit (Sigma) according to manufacturer’s instructions.

Bgl2 accumulation was determined as described (Curwin et al., 2012) with minor modifications. Cells were grown in YPR to log phase, transferred to YPG for 2 h, and cells were either shifted to 37 °C (*sec14-1*) or incubated with 0.5 mM NAA (*boi1Δ boi2-aid*). After 2 hours, equal cell numbers were harvested by centrifugation and washed in 10 mM NaN_3_/NaF solution. Total protein extracts were prepared by TCA extraction. For the internal fraction, cells were resuspended in pre-spheroplasting buffer (100 mM Tris- H_2_SO_4_, pH 9.4; 50 mM β-mercaptoethanol; 10 mM NaN_3_; 10 mM NaF) and washed with spheroplasting buffer without zymolyase (50 mM KH_2_PO_4_-KOH, pH 7.4; 1.4 M sorbitol; 10 mM NaN_3_). Then, cells were resuspended in spheroplasting buffer containing 167 μg/ml zymolyase 100T (Seikagaku Biobusiness), and incubated with gentle mixing. Spheroplasts were harvested and resuspended in sample buffer before separation by SDS–PAGE and western blotting to detect Bgl2 (specific antibody gift from Randy Schekman, University of California, Berkeley) and Glucose-6-phosphate dehydrogenase as a loading control (Sigma).

## Acknowledgements

We are grateful to Karolina Jodkowska for assistance with the initial characterization of *EXO70*\*. We thank Yves Barral, Amy Curwin, Vivek Malhotra, Snezhana Oliferenko, Isabelle Sagot, Jerome Solon, and members of the Mendoza lab for helpful suggestions; Marti Aldea, Eric Bailly, Derek McCusker and Randy Schekman for providing reagents; the CRG Genomics and Advanced Light Microscopy Units; and the CCiT Electron Cryomicroscopy Unit (Universitat de Barcelona). This research is supported by grants from the Spanish ministry of Economy and Competitiveness (BFU09-08213 and BFU2012-37162 to MM, and BIO2012-37161 to TG), the Qatar National Research Fund (NPRP 5-298-3-086 to TG), and the European Research Council (ERC Starting Grant 260965 to MM and 310325 to TG). LPP was funded through La Caixa - CRG International Fellowship Program.

The authors declare no conflicting financial interests.

## SUPPLEMENTARY LEGENDS S1 to S2

### SUPPLEMENTARY TABLES S1 to S4

**Fig. S1. Genetic suppression in a *boi1Δ boi2Δ* strain.** (A) Tetrad analysis of *boi1Δ BOI2 / BOI1 boi2Δ* diploid (product of a cross between *boi1Δ* and *boi2Δ* haploids) grown for 3 days at 25 °C. (B) Tetrad analysis of *boi1Δ boi2Δ SUP / BOI1 BOI2* diploid (from a cross between *boi1Δ boi2Δ SUP* [from Norden et al., 2004] and *BOI1 BOI2*) grown for 3 days at 25 °C. (C) Serial dilutions of the indicated strains spotted onto the indicated plates and incubated for 5 days at 30 °C. Clones 1-3 are derived from independent spores obtained from a cross between *boi1Δ* and *boi2Δ EXO70*\*.

**Fig. S2. Depth of coverage analysis of wild type and *boi1Δ boi2Δ*.** Chromosomes of wild type (*WT*, top) and *boi1Δ boi2Δ SUP* mutant (bottom) were analyzed by whole genome sequencing depth of coverage. Chromosomes are given on the X-axis, while read density is given on Y-axis. Duplications are visible as peaks, i.e. the rDNA cluster is expanded in both strains. We found no difference in copy number between these strains, except for *URA3* used to delete *BOI2* and present in approximately 10 copies, visible as a peak in chromosome V. Two copies of the *LEU2* marker used to delete *BOI1*, and deletions of *BOI1* and *BOI2* were also detected but are not visible at the scale shown.

**Fig. S3.** Internal and total Bgl2 levels determined by western blot, of cells of the indicated strains after incubation with DMSO or NAA for 2 h (*boi1Δ boi2-aid*), or after shift to 37 °C for 2 h (*sec 14-1*). G6PDH was used as a loading control.

**Fig. S4.** Correlation between fluorescence intensity levels of Sec4-GFP at the bud neck and in the whole cell, measured in sum projections of confocal z-stacks spanning the entire cell.

**Table S1.** Coding, non-synonimous SNPs found in *boi1Δ boi2Δ* cells compared to its parental wild-type. The table lists the chromosome and position of the SNP, the affected gene and codon in wild type and mutant strains, the predicted aminoacid change and the reported gene function according to http://www.yeastgenome.org.

| Location | Name | WT | *boi1Δ*<br>*boi2Δ* | aa<br>change | Gene function |
| --- | --- | --- | --- | --- | --- |
| chrII:39982 | *BNA4* | GAT | AAT | D281N | biosynthesis of NAD |
| chrIII:59865 | *GFD2* | ATA | ATG | I280M | unknown |
| chrV:221858 | *YER034W* | ACC | AGC | T5S | unknown |
| chrVIII:186001 | *MSC7* | ATT | GTT | I268V | unknown |
| chrX:273982 | *EXO70* | GGA | AGA | G388R | Subunit of the exocyst complex |
| chrXII:799068 | *NUP2* | TCG | CCG | S547P | Nucleoporin involved in nucleocytoplasmic transport |
| chrXVI:390476 | *YPL085W* | GCG | GAG | A1138E | COPII vesicle coat protein required for ER transport vesicle budding |

**Table S2.**
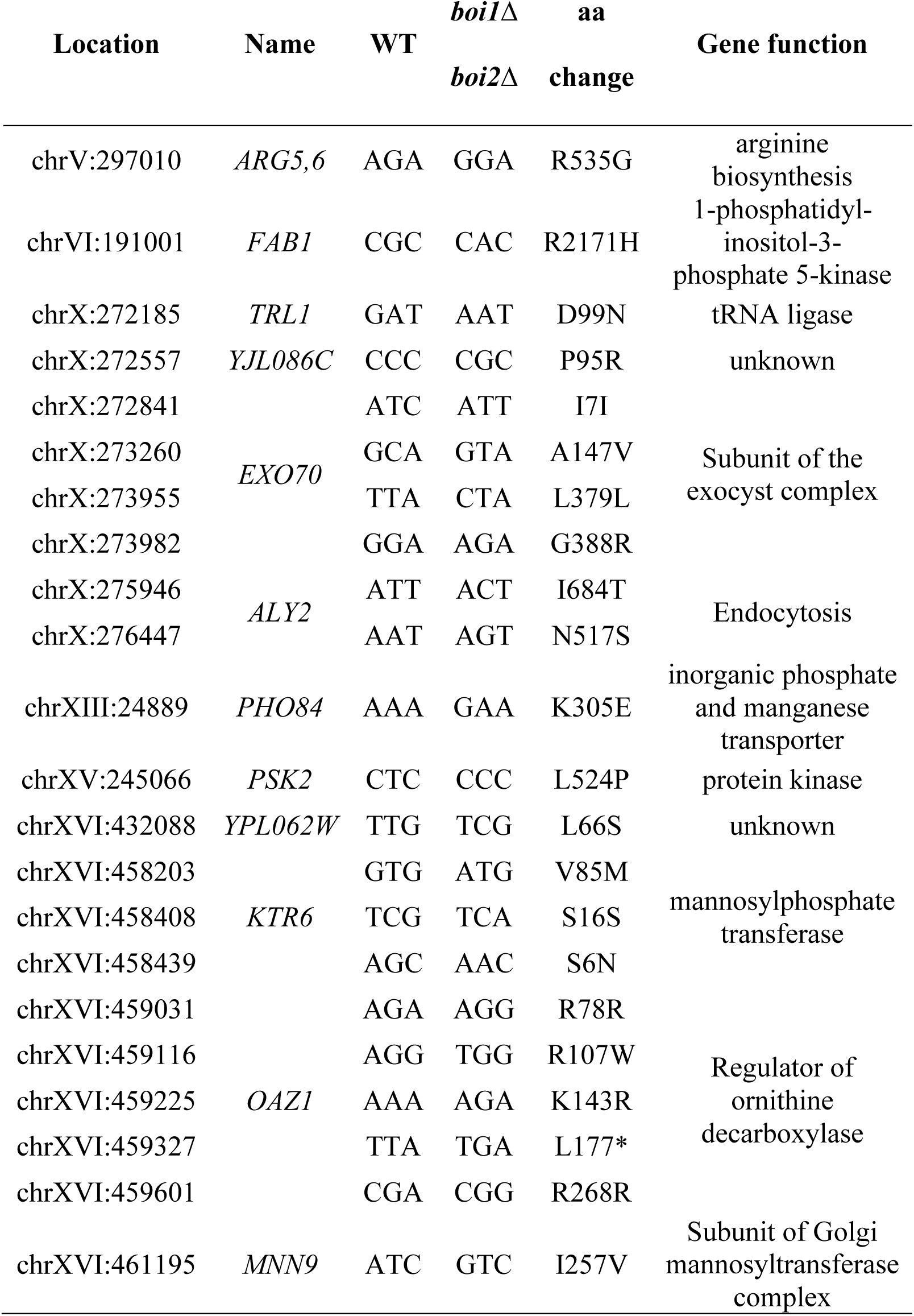
Non-synonymous SNPs present in two meiotic products of a *boi1Δ/BOI1 boi2Δ/BOI2 SUP*/+ zygote (*boi1Δ boi2Δ*) and absent in the remaining two (*BOI1+ BOI2+*), as well as absent from the wild type. Assembly statistics for *de novo* assembled genomes are listed in table S3.

**Table S3.**
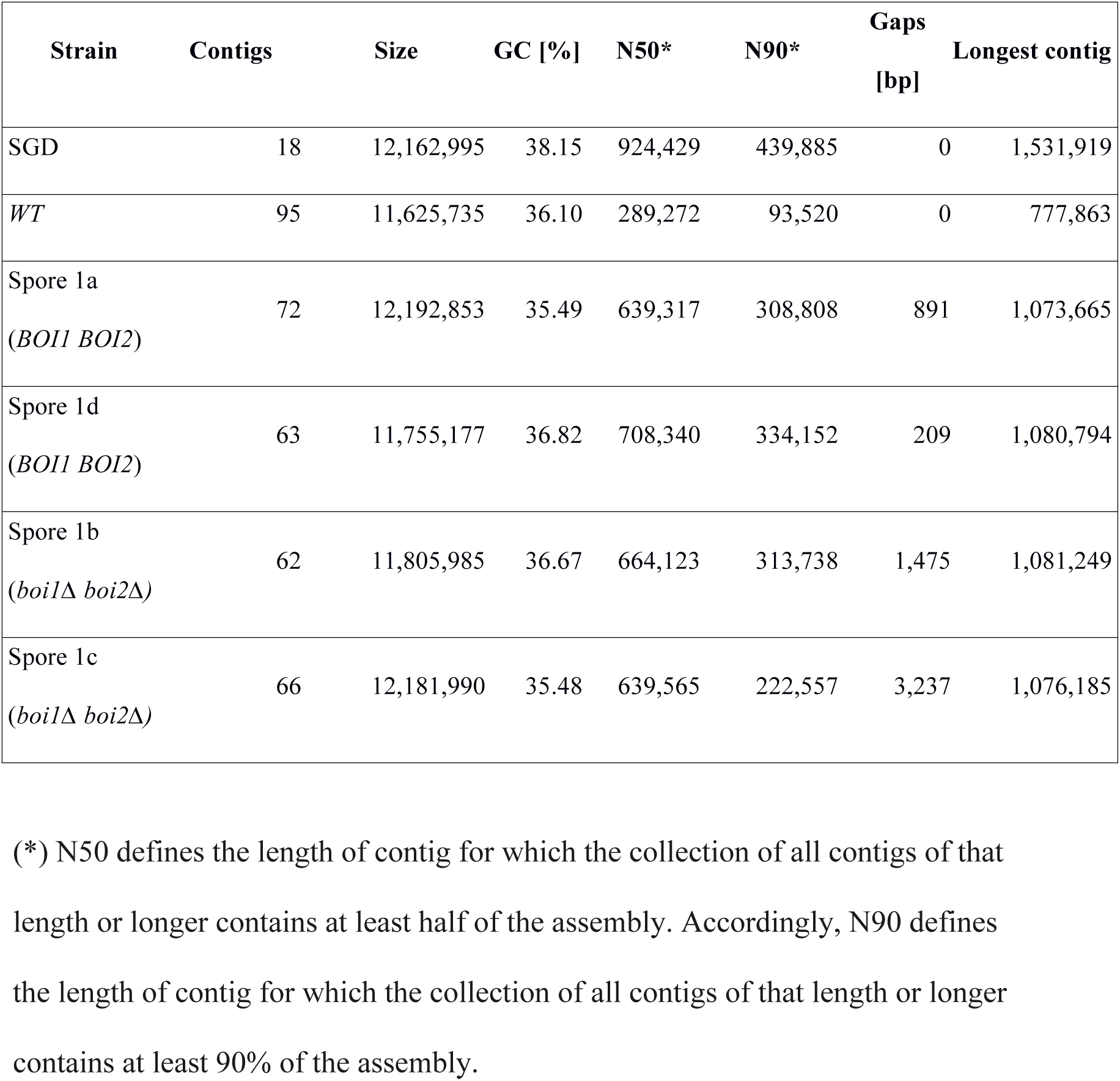
Genome assembly statistics for *de novo* assembled genomes used to generate table S2. The table lists: strain name, number of contigs, cumulative assembly size, GC content, N50, N90, size of gaps and the longest contig length are given. *S. cerevisiae* reference genome (SGD) statistics are given at the top.

| Strain | Contigs | Size | GC [%] | N50* | N90* | Gaps<br>[bp] | Longest contig |
| --- | --- | --- | --- | --- | --- | --- | --- |
| SGD | 18 | 12,162,995 | 38.15 | 924,429 | 439,885 | 0 | 1,531,919 |
| *WT* | 95 | 11,625,735 | 36.10 | 289,272 | 93,520 | 0 | 777,863 |
| Spore 1a<br>(*BOI1 BOI2*) | 72 | 12,192,853 | 35.49 | 639,317 | 308,808 | 891 | 1,073,665 |
| Spore 1d<br>(*BOI1 BOI2*) | 63 | 11,755,177 | 36.82 | 708,340 | 334,152 | 209 | 1,080,794 |
| Spore 1b<br>(*boi1Δ boi2Δ*) | 62 | 11,805,985 | 36.67 | 664,123 | 313,738 | 1,475 | 1,081,249 |
| Spore 1c<br>(*boi1Δ boi2Δ*) | 66 | 12,181,990 | 35.48 | 639,565 | 222,557 | 3,237 | 1,076,185 |

(\*) N50 defines the length of contig for which the collection of all contigs of that length or longer contains at least half of the assembly. Accordingly, N90 defines the length of contig for which the collection of all contigs of that length or longer contains at least 90% of the assembly.

